# Microbial community dispersal in sourdough

**DOI:** 10.1101/2021.10.18.464797

**Authors:** Lucas von Gastrow, Rémy Amelot, Diego Segond, Stéphane Guézennec, Florence Valence, Delphine Sicard

## Abstract

Understanding how microbes disperse in ecosystems is critical to understand the dynamics and evolution of microbial communities. However, microbial dispersal is difficult to study because of uncertainty about the vectors that may contribute to their migration. This applies to both microbial communities in natural and human-associated environments. Here, we studied microbial dispersal among French sourdoughs and flours used to make bread. Sourdough is a naturally fermented mixture of flour and water. It hosts a community of bacteria and yeasts whose origins are only partially known. We analyzed whether flour is a carrier of sourdough yeast and bacteria and studied whether microbial migration occurs between sourdoughs. The microbial community of a collection of 46 sourdough samples, as well as that of the flour from which each was made, was studied by 16S rDNA and ITS1 metabarcoding. No sourdough yeast species were detected in the flours. Sourdough lactic acid bacteria (LAB) were found in only five flour samples, and they did not have the same amplicon sequence variant (ASV) as found in the corresponding sourdough. The species shared between the sourdough and flour samples are commonly found on plants and are not known to be alive in sourdough. Thus, the flour microorganisms did not appear to grow in the sourdough microbial community. Dispersal between sourdoughs was also studied. Sourdoughs shared no yeast ASV, except in few cases where groups of three to five bakers shared some. These results suggest that there is little migration between sourdoughs, except in a few situations where bakers may exchange sourdough or be vectors of yeast dispersal themselves.

## 1 Introduction

Understanding the functioning and evolution of communities is central to ecological studies. Many of the concepts and debates that have animated this field have arisen from the study of plant communities (Mikkelson, 2005). Microbial communities have also been a subject of increasing interest, and it is now clearly established that they play a central role in the functioning and evolution of many ecosystems. Numerous concepts have been proposed in community ecology but it is only recently that theoretical models have unified them to take account of local evolutionary dynamics and the links between communities. Vellend (2010) defined four factors that shape communities : diversification, selection, dispersal and drift, and more recently, Thompson et al. (2019) proposed a meta-community model with three factors : density-independent responses to abiotic conditions, density-dependent biotic interactions, and dispersal. These general frameworks offer valuable tools to understand the dynamics of microbial communities but suffer from a lack of empirical data on the selection processes and dispersal of microbial communities.

Microbial communities are present in both wild environments and in all human-associated environments. They have been used to make fermented foods since the Neolithic era (Tamang and Kailasapathy, 2010), in which they usually display relatively little complexity with regards wild environments, making them good model systems for ecological studies. They are organized as metacommunities in which the microbial community of each leaven evolves as a function of human practices and may be linked to others through exchanges of the leavens themselves or of the raw materials used to feed them. Among these numerous fermented foods, sourdough microbial communities used for bread-making represent a good metacommunity model system. First, sourdough microbial communities are relatively simple, usually containing one to two dominant bacterial and yeast species (Carbonetto et al., 2018; Arora et al., 2021). Second, sourdoughs are made of few ingredients, basically flour and water, which are regularly added to feed the microorganisms, thus limiting the number of sources of microbial species. Third, the microbial communities in sourdough have been widely studied and reviewed (De Vuyst et al., 2016; Gänzle and Ripari, 2016; Gobbetti et al., 2016; Gänzle and Zheng, 2019; Arora et al., 2021; Van Kerrebroeck et al., 2017; Calvert et al., 2021; Lau et al., 2021). Well known species such as *Fructilactobacillus sanfranciscensis, Lactiplantibacillus plantarum, Levilactobacillus brevis* bacteria and *Saccharomyces cerevisiae, Kazachstania humilis, Torulaspora delbrueckii* and *Wickerhamomyces anomalus* yeasts are frequently encountered. Finally, population genomic analysis of the yeast species *S. cerevisiae* has shown that sourdough yeast populations differ from commercial yeasts and may have undergone specific selection processes when compared to industrial processes (Bigey et al., 2020).

Although the microbial composition of sourdough has been well described, the origins and dispersal of sourdough microbial species have only been partially studied. Previous studies showed that the same species of lactic acid bacteria (LAB) or yeast could be found on the baker’s tools (Minervini et al., 2015) or hands (Reese et al., 2020) and in their sourdough. But this does not tell us whether the microorganisms in the sourdough came from the baker’s tools or hands or vice versa. Moreover, no sourdough microorganisms were detected in the bakery air (Minervini et al., 2015) or in the water (Scheirlinck et al., 2009; Reese et al., 2020) used to make the sourdough. Finally, other studies have shown that flour can be a vector for *Lactobacillaceae*. However, this was only shown for three different flours (Minervini et al., 2018a) or for laboratory-made sourdoughs (De Angelis et al., 2019), whose dynamics are not the same as bakery sourdoughs (Minervini et al., 2012). The source of sourdough yeast and bacteria therefore still needs to be elucidated.

In France, analyses of sourdough microbial communities revealed that *F. sanfrancisensis* was the dominant bacterial species in almost all sourdoughs (Lhomme et al., 2015; Michel et al., 2016). Yeast species were more diverse and included *S. cerevisiae* but also many different *Kazachstania* species Urien et al. (2019). The distribution of the latter was associated with the type of bread-making practices. Sourdough made by farmer bakers tended to carry *K. bulderi* while sourdough made according to artisanal practices often contained *K. humilis* (Michel et al., 2019). While farmer bakers exchange seeds, share mills or supply each other with flour, artisanal bakers usually buy their flour from millers who produce and store flour at a larger scale. Different sources of flour supply may lead to different pathways for microorganism dispersal and explain the structuring of yeast species diversity as a function of bread-making practices.

To test this hypothesis, we analyzed the role of flour in the dispersal of sourdough microor-ganisms among French bakers and farmer-bakers. We studied the microbial species diversity of 46 flours and related sourdough samples as well as the bread-making practices of the bakers. We did not find any evidence that flour is a vector for sourdough yeasts. Flour can carry LAB species but these are not the same as those found in mature sourdough, suggesting another origin for sourdough LABs. We also studied whether microbial dispersal occurred between sourdoughs. We found that sourdough shared the same LAB ASVs but most of them have their own yeast ASVs composition suggesting that there is little exchange between sourdoughs.

## 2 Material and Methods

### 2.1 Survey of bread-making practices

A total of 22 bakers and 22 French farmer-bakers completed a questionnaire on their bread-making practices, as described by Michel et al. (2019). Questions concerned sourdough management (addition of bran, back-slopping technique, time elapsing since sourdough initiation, sourdough hydration, number of back-slopping procedures per week and between bread-making sessions, temperature at back-slopping), the flour (self-produced or not, type of cereal variety, type of mill) and the bread-making process (use of selected baker’s yeast in bread or in other products, mechanical or manual kneading, proportions of sourdough, flour, water and salt in bread dough, fermentation time, quantity of bread produced each week, number of bread-making sessions per week). We also asked the producers if they had shared raw materials (grains, flour or sourdough) or if they had physical contacts with each other.

### 2.2 Sample collection

A total of 46 sourdoughs were collected, together with the flour used to make each one. Forty-four sourdoughs came from different bakeries, and two bakeries (B64 and B68) sent two sourdoughs, so that 46 sourdough and 44 flour samples were studied. Samples were collected between September 2018 and July 2019 and were received at the laboratory within one to three days. All samples were stored at −20°C in plastic bags and plastic tubes, respectively, before DNA extraction.

### 2.3 Sourdough and flour microbial enumeration and strain isolation

All 46 sourdoughs and 21 flour samples were plated at reception. 10 g sourdough or 3 g flour were diluted ten times in tryptone-salt buffer (1 g/L tryptone, 8 g/L NaCl). After serial dilutions, lactic acid bacteria (LAB) were enumerated on MRS-5 (Meroth et al., 2003) with 100 μg cycloheximide and on PCA (6 g/L Tryptone, 2.5 g/L yeast extract, 1 g/L glucose, 15 g/L agar) media while yeasts were enumerated on YEPD medium (10 g/L yeast extract, 20 g/L peptone, 20 g/L dextrose, 100 mg/L chloramphenicol).

### 2.4 Identification of bacterial and yeast species

The species diversity of the sourdoughs and flours was analyzed by amplicon-based DNA metabarcoding using two separate Illumina MiSeq runs to prevent any contamination between sample types.

#### 2.4.1 DNA extraction from sourdough and flour

DNA was extracted using a Qiagen PowerSoil DNA isolation kit (12888-100). Sourdough DNA was extracted directly from 200 mg of material following the kit procedure. For the flour, 3g of material was washed in sterile PBS, filtered in a sterile filter bag (BagPage+, Interscience, France) and concentrated in 500 μL PBS after centrifugation. Extraction was then performed in accordance with the manufacturer’s instructions.

#### 2.4.2 MiSeq sequencing

The 16S V3-V4 region was amplified for bacteria and the ITS1 region for fungi. For fungi, the ITS1 region was targeted with the PCR primers ITS1-F (5’ - CTTGGTCATTTAGAGGAAGTAA - 3’) and ITS2 (5’ - GCTGCGTTCTTCATCGATGC - 3’) (White et al., 1990), while for bacteria, the 16V3-V4 region was targeted with the PCR primers 343F : (5’ - TACGGRAGGCAGCAG - 3’) and 784R : (5’ - TACCAGGGTATCTAATCCT - 3’) (Liu et al., 2007). The primers also included the Illumina tail (5’ - TCGTCGGCAGCGTCAGATGTGTATAAGAGACAG - 3’), and a frame-shift of four, six or eight random nucleotides for forward primers and four, five or six random nucleotides for reverse primers, in order to prevent saturation during sequencing. The resulting primers therefore had the following structure : 5’ - Illumina tail - frame-shift - genome targeting region - 3’. All the primers used are listed in Supplementary Materials Table S1. For each forward or reverse primer, an equimolar mix of the three primers containing the different frame-shifts was added to the PCR mix. To prepare the multiplexed Illumina libraries, we employed a strategy based on a two-step PCR approach : a first PCR using the locus-specific primers including the Illumina adapter overhang (with 30 cycles), and a second PCR for the incorporation of Illumina dual-indexed adapters (with 12 cycles). Bead purifications were carried out after each PCR. Quantification, normalization and pooling were performed before sequencing on Illumina MiSeq (Ravi et al., 2018).

#### 2.4.3 Bioinformatics analyses

The resulting sequences were analyzed using R (Team, 2019) workflow combining dada2 v.1.16 (Callahan et al., 2016) and FROGS 3.1.0 (Escudié et al., 2018) software. Reads were filtered, merged and assigned to ASVs with dada2 and the ASVs were assigned to species using the FROGS affiliation tool. Adapters were first removed using cutadapt v. 1.12 with Python 2.7.13. Reads were then filtered using the dada2 filterAndTrim function, with a truncation length of 250 bp for ITS1 forward and reverse reads and 275 and 200 bp for 16S forward and reverse reads, respectively. This truncation reduced the error rate while still allowing the merging of most reads. The error model was then calculated using the learnErrors function. Reads were dereplicated using derepFastq and the dada2 core sample inference algorithm was executed. Forward and reverse reads were then merged with a minimum overlap of 20 bp. The resulting sequences were saved in a sequence table using makeSequenceTable. Chimera were removed using the removeBimeraDenovo function. The amplicon sequence variants (ASV) in the sequence table were then assigned to species using FROGS affiliation v3.2.2 with silva 138 (Quast et al., 2013) for 16S and Unite 8.0 (Nilsson et al., 2019) for ITS1. Unite was completed with ITS1 reference sequences from yeast species usually found in sourdough. Multi-affiliations were dealt with by assigning the lowest common taxonomy level to multi-affiliated ASVs. Samples were rarefied to the minimum number of reads for each barcode, or 1000 reads using the rarefy_even_depth function of the R (v. 4.1.0) phyloseq package (v. 1.24.2) (McMurdie and Holmes, 2013). Samples with a depth of less than 1000 were discarded. If not otherwise specified, the analyses were conducted on the rarefied data.

#### 2.4.4 Analysis of bread-making practices

Groups of bread-making practices were obtained with an MCA computed with the R package FactoMineR (v. 2.4), and individuals were clustered using the HCPC function with two clusters. They were plotted using the factoextra package (v. 1.0.7).

#### 2.4.5 Statistical analysis

A Wilcoxon-Mann-Witney test was performed to compare the diversity index between the flour and sourdough samples. The correlation between flour and sourdough diversity was computed using a Spearman rank-order correlation test. Both tests were computed using the R package stats v 3.6.2, with the wilcox.test and cor.test functions, respectively. A Mantel test was performed to test the link between geographical distances for sourdoughs and Bray-Curtis distance matrices, using the R ape package (v. 5.5) mantel.test function.

## 3 Results

### 3.1 The sourdough microbiota had greater microbial density but less species diversity than the flour microbiota

To analyze the role of flour microbiota in the composition of sourdough microbiota, we compared 46 sourdough samples obtained from 44 bakeries with the 44 flour samples used to refresh them (only 21 flour samples were plated for microbial counts).

On average, microbial density was higher in sourdoughs than in flours, for both bacteria and fungi. Sourdoughs contained on average 1.9 ∗ 10^7^ (sd = 1.3 ∗ 10^7^) CFU/g (colony forming units/g) of yeast while flours contained a mean of 2.3 ∗ 10^3^ (sd = 1.6 ∗ 10^3^) CFU/g. As for bacteria, the sourdoughs contained 1.3 ∗ 10^9^ (sd = 1.3 ∗ 10^9^) CFU/g while flours contained 7.7 ∗ 10^3^ (sd = 2.0 ∗ 10^4^) CFU/g or 6.9 ∗ 10^4^ (sd = 1.0 ∗ 10^5^) CFU/g, depending on whether the estimation of bacterial density was performed on MRS or PCA. Sourdoughs were only plated on MRS medium, as we expected to find only *Lactobacillaceae*, while flour generally harbors a more diverse bacterial community so we also plated these samples on PCA, which is a less specific medium. The observation of fungal morphology on YEPD petri dishes revealed that most flour samples contained filamentous fungi, some with a typical *Penicillium* morphology, while sourdough samples were characterized by the presence of yeasts.

Although sourdoughs had a higher microbial density than flour, their microbial communities were less diverse than those in flour. Alpha diversity indexes calculated on the number of bacterial and fungal species were significantly lower in sourdough than in flour in terms of both richness (Wilcoxon-Mann-Witney test, bacteria W = 1725.5, *P <* 0.001, fungi W = 1555.5, *P <* 0001) and evenness (Wilcoxon-Mann-Witney test, bacteria W = 1929, *P <* 0.001, fungi W = 1467, *P <* 0001 ; Figure 1). This difference was greater for bacteria than for fungi, with averages of four and 11 species for bacteria in sourdough and flour, respectively, and 10 and 13 species for fungi in sourdough and flour, respectively.

**Figure 1-.**
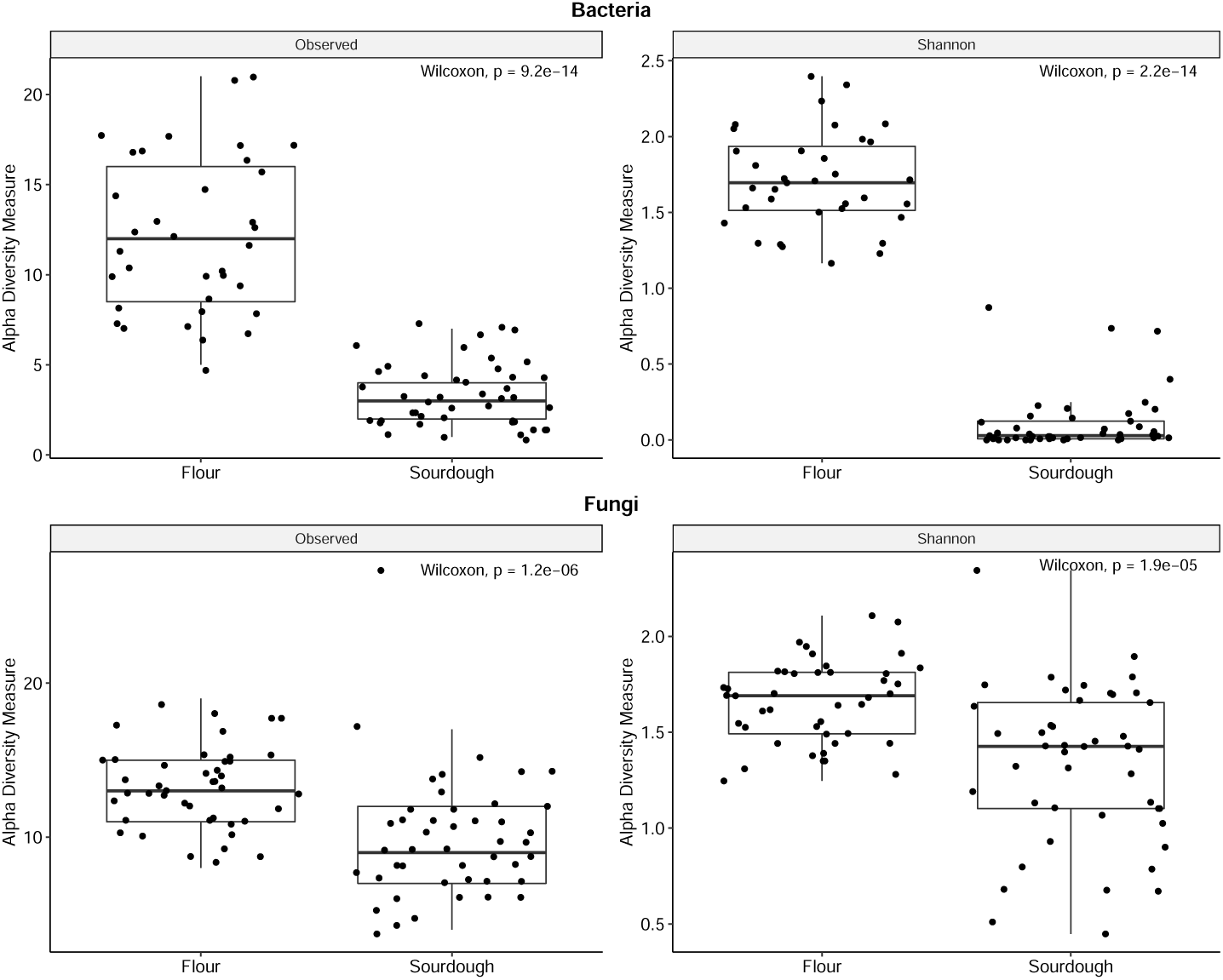
Alpha diversity in sourdough and flour samples, estimated from 16S V3-V4 and ITS1 Illumina MiSeq reads assigned to species. Species richness (on the left) and evenness (on the right) are plotted.

Sourdough species diversity was not correlated with flour species diversity for either bacteria (Spearman = 13617, P = 0.86) or fungi (Spearman = 13019, P = 0.91).

The microbiota compositions of sourdough and flour were characterized by different families. The bacteria in the sourdoughs were almost entirely composed of *Lactobacillaceae*, while flour contained mainly *Erwiniaceae* and *Pseudomonadaceae*. In sourdough, all samples but three contained *Fructilactobacillus sanfranciscensis* as the dominant bacterial species ; the others contained *Companilactobacillus paralimentarius*. Less frequently, the presence of *Levilactobacillus brevis, Latilactobacillus* sp. and *Lactilactobacillus* sp. was found. In flour, *Erwiniaceae, Pantoea agglomerans*, an unidentified *Pantoae* sp., and *Pseudomonadaceae* were generally detected. Among *Pseudomonas* sp., some were *P. graminis, P. rhizospherae* or *P. donghuensis*. As for fungi, *Saccharomycetaceae* was determined in most sourdough samples but was almost absent from flour samples (Figure 2) ; *S. cerevisiae* was found in 14 sourdough samples, *K. humilis* in seven samples and *K. bulderi* in six. These species were never found in flours. *Pleosporaceae* species (*Alternaria alternata* and *Alternaria infectoria*), *Mycosphaerellaceae* (*Mycosphaerella tassiana*) and an unidentified fungus from the *Dothideomycetes* family were detected at a high frequency in almost all flour samples.

**Figure 2-.**
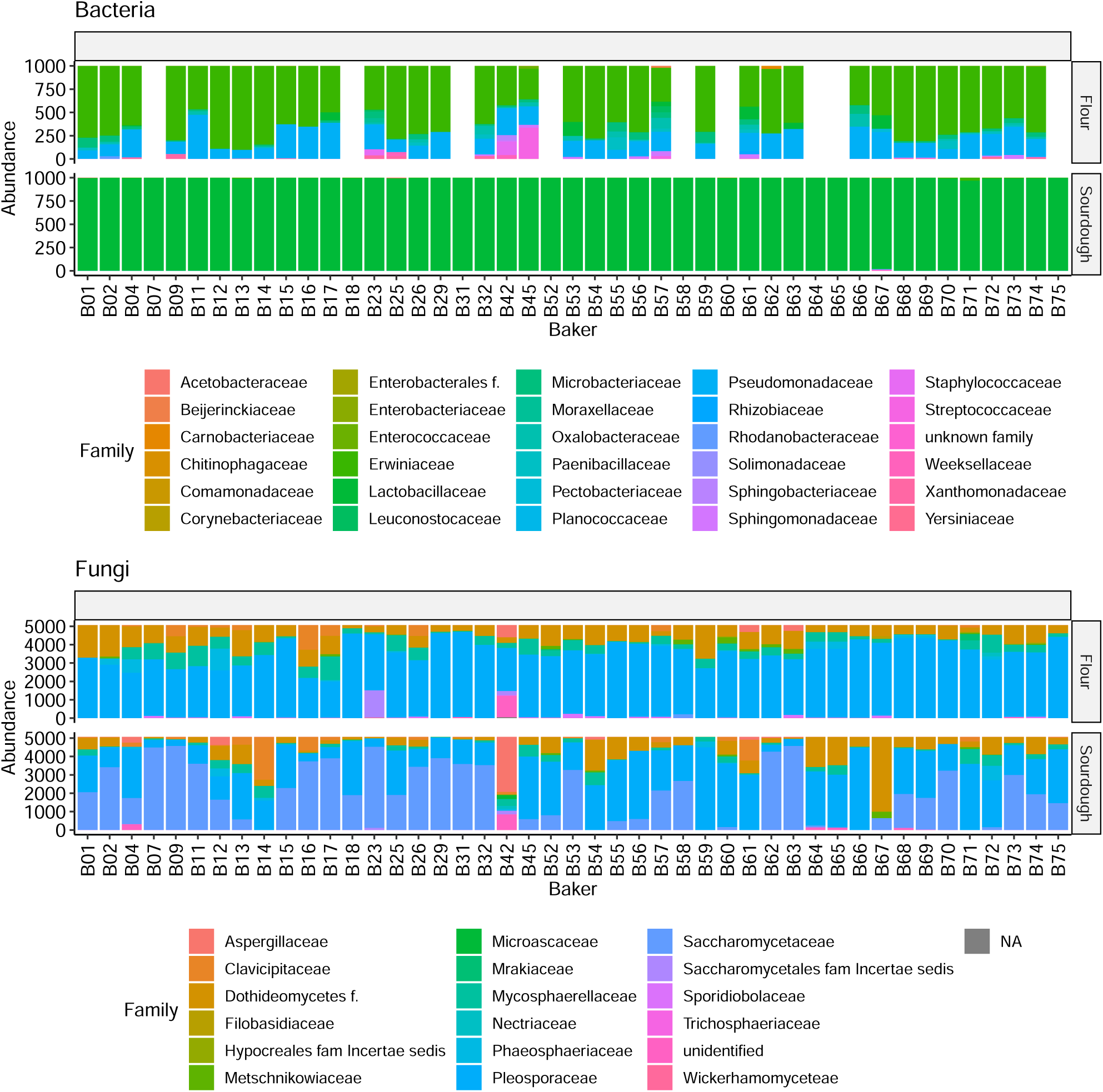
Abundance of the different families in flours and sourdoughs. White bars represent the different ASVs.

### 3.2 Very little overlap between the microbiotas of sourdough and flour

Any overlaps between the sourdough and flour communities were analyzed using the Weighted Bray-Curtis distance calculated on the basis of species diversity. The Weighted Bray-Curtis was used to build two PCoAs, one for the bacterial community and the other for the fungal community. PCoA axis 1 and 2 explained 79.1% and 8.5% of variance for bacteria, and 28.5% and 13.6% of variance for fungi (Figure 3). For bacteria, axis 1 separated the flour and sourdough bacterial communities. For fungi, axis 1 separated many but not all of sourdough fungal communities from flour communities. Over the 46 sourdough fungal communities, 14 co-localized with flour fungal communities. Flour and sourdough dissimilarity matrices were not correlated (Mantel test, z = 836, p = 0.667 for bacteria and z = 854, p = 0.13 for fungi). Close microbial communities among flours did not lead to close microbial communities among sourdoughs.

**Figure 3-.**
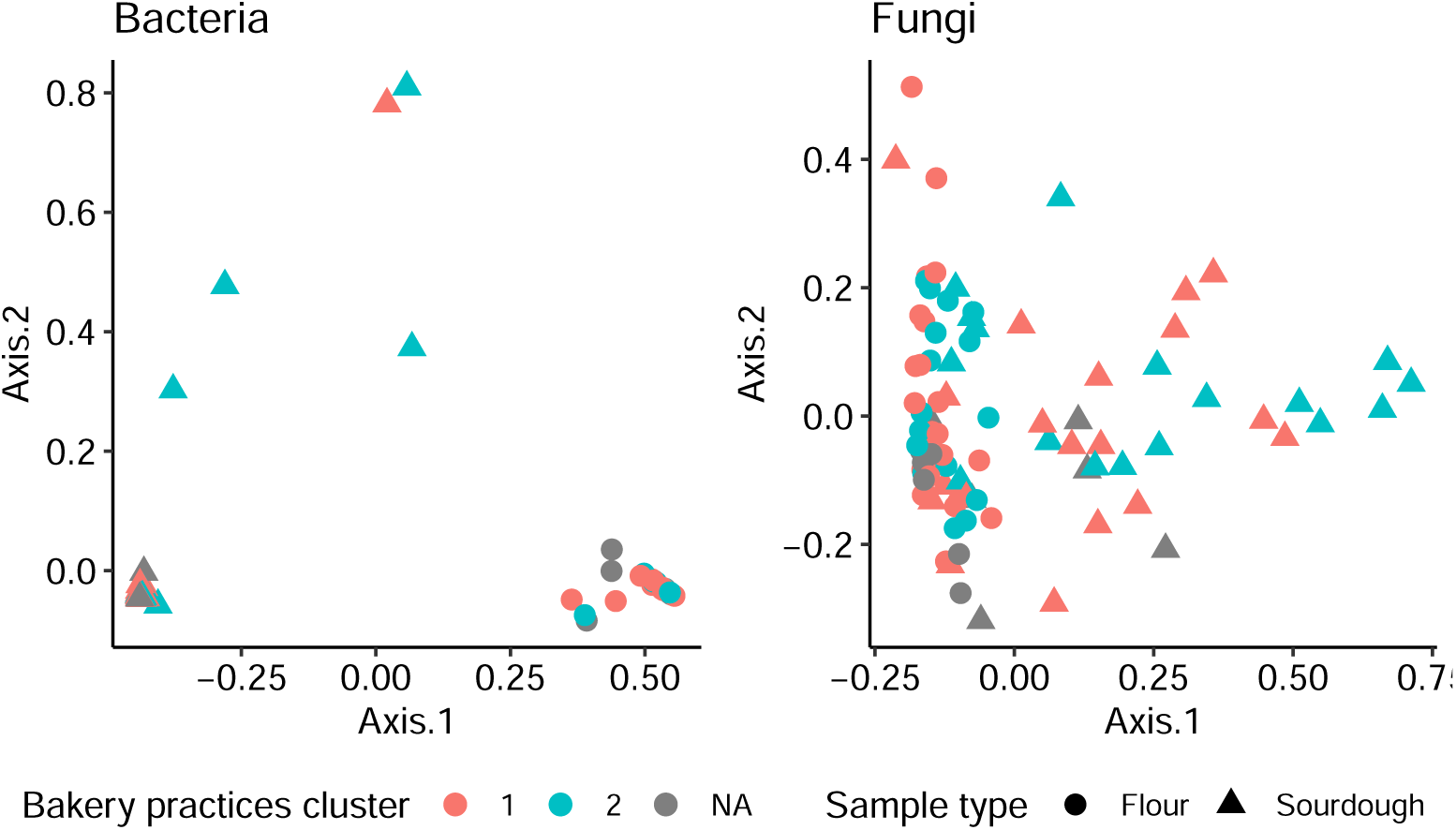
PCoA based on Bray-Curtis dissimilarity for bacteria (left) and fungi (right). Bray-Curtis dissimilarity was computed on the basis of the abundance of the different species. Each point represents a sample. Colors indicate the bakery practices cluster, with farmer-baker practices in red and artisan-baker practices in blue. Sample types are represented by different shapes, flours being shown as circles and sourdoughs as triangles.

We analyzed bread-making practices in order to determine whether they might be related to microbial communities in sourdough and flour. Two groups of bread-making practices could be distinguished (Figure S1). Farmer-baker practices (cluster 1) were more frequently associated with the use of non-commercial yeast, ancient wheat landraces, small production runs and lengthy fermentation while artisanal practices (cluster 2) were generally characterized by larger scale production, short fermentation, and the use of commercial yeast and modern wheat varieties. Sourdough from farmer-bakers frequently contained *K. bulderi* as the dominant yeast species. However, analysis of the association between sourdough and flour microbial community dissimilarity and the geographical distances between bread-making practices did not reveal any correlation (Mantel test, for flour, z = 308, p = 0.59 and z = 235, p = 0.79 for bacteria and fungi, respectively ; for sourdough, z = 153, p = 0.60 and z = 411, p = 0.32 for bacteria and fungi, respectively).

The differences between the microbial communities in sourdough and flour were explained by the high abundance in sourdough samples of fermentative microorganisms, which were almost never found in the flour samples. (Figure 4).

**Figure 4-.**
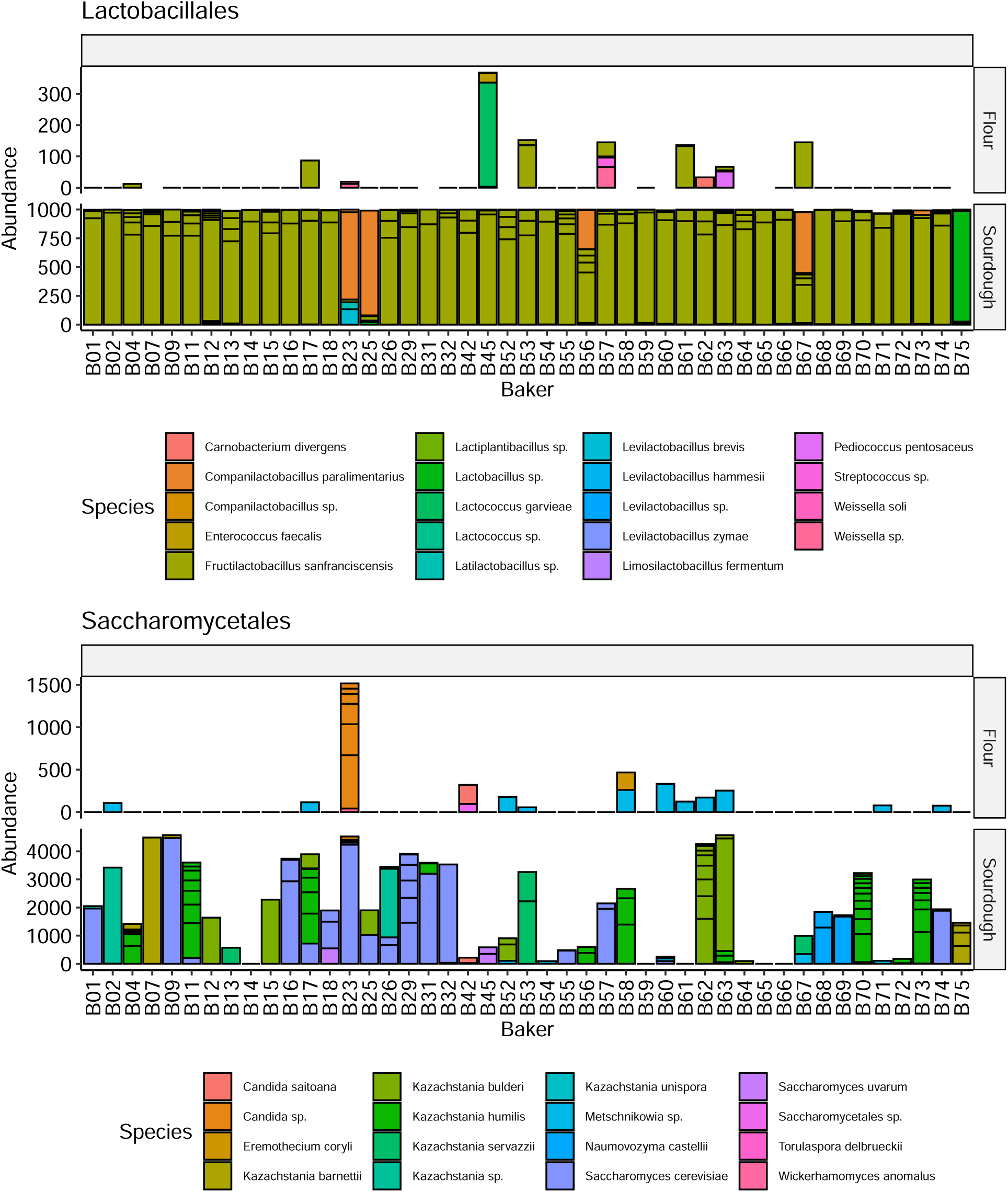
Abundance of *Lactobacillales* and *Saccharomycetales* in flour and sourdough. The axes have different scales for abundance in flour and sourdough.

Overall, fermentative bacteria in the *Lactobacillales* order and yeast in the *Saccharomycetales* order were not detected in most flour samples. Out of 46 samples, ten flour samples contained fermentative bacterial species (*F. sanfranciscensis, Lactococcus garviae, Carnobacterium divergens, Weisella* or *Streptococcus* species) and 13 harbored fermentative yeasts (*Candida saitoana*, an unidentified *Candida* species, *Wickerhamomyces anomalus, Mechnikovia* sp. or *Eremothecium coryli*). However, the fermentative species found in flour samples were generally not found in the related sourdoughs. In six cases, *F. sanfranciscensis* was found in both flour and sourdough. Nevertheless, in these cases, the ASVs were not the same except in the case of baker 53 (Figure 5). *Lactococcus garviae* was found in the flour and sourdough used by baker 45 but only one read was present in the sourdough and this ASV differed from that found in the flour. An unidentified *Metschnikowia* species was found in four pairs of sourdough and flour, and *Candida saitoana* and an unidentified *Candida* species in one pair of sourdough and flour samples, although the same ASV was not found in them. Many non-fermentative fungal species were shared between flour and sourdough samples. They were mainly filamentous fungi, and notably species from the genus *Alternaria* or *Mycosphaerella*. For these species, the flour and sourdough samples shared on average 0.98 ASV (sd = 1.48).

**Figure 5-.**
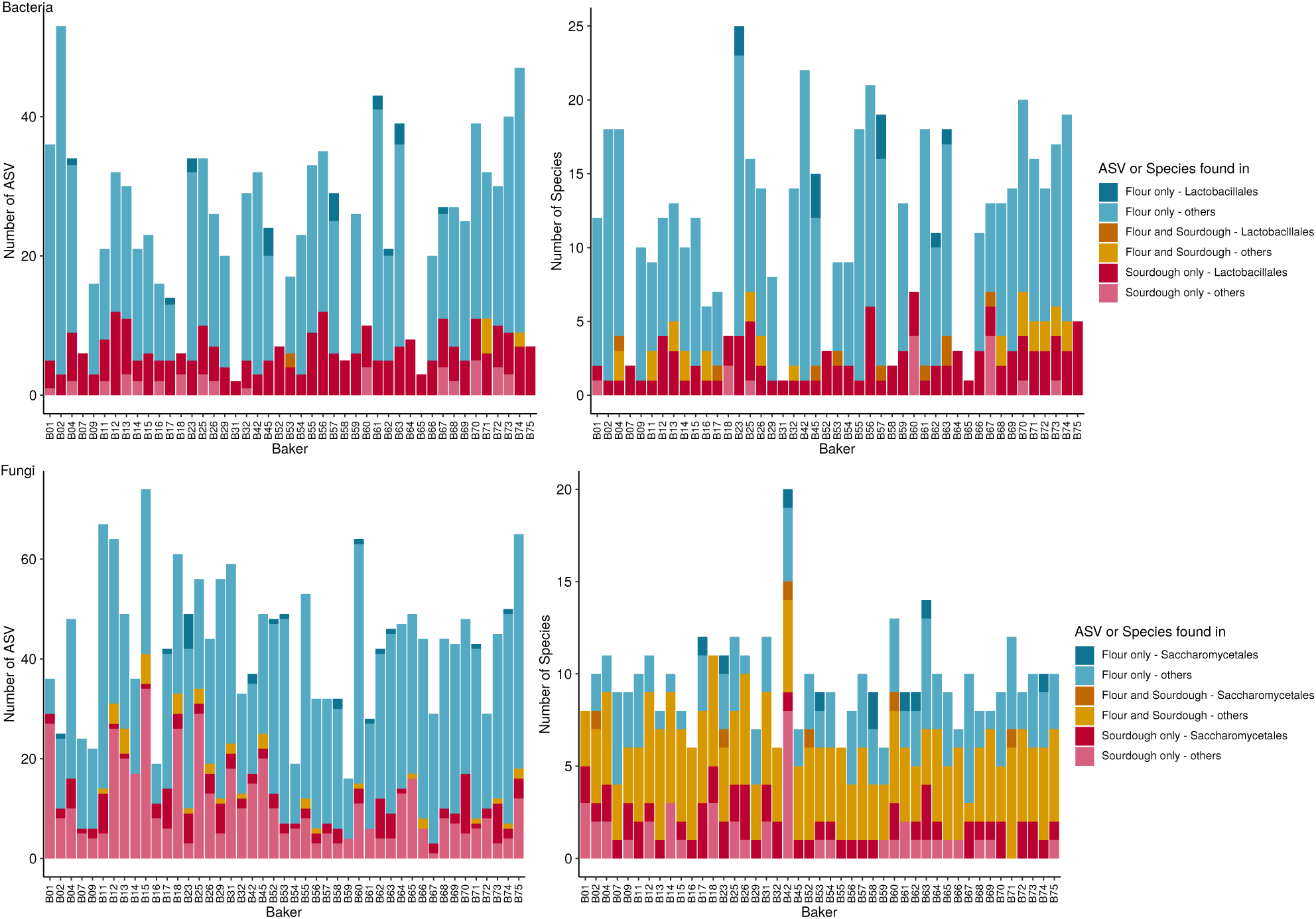
Number of shared species (on the right) and ASV (on the left) between sourdoughs and the flour used to make them. Results for bacteria are shown at the top and for fungi at the bottom.

### 3.3 Dispersal appear to be reduced between sourdoughs

The poor overlap between the microbiotas of flours and sourdoughs suggested that flour is not a vector of microbial dispersion between bakeries. However, microbial dispersal could occur through direct exchanges of sourdough between bakers. We therefore analyzed the microbial flux between sourdoughs by looking at the number of sourdoughs containing the same ASVs ; those of the *F. sanfranciscensis* species were shared on average by 5.8 sourdoughs (sd = 9.4), while ASVs from the *Saccharomycetales* yeasts were shared by 1.22 sourdoughs on average (sd = 0.69).

We then studied the occurrence of ASVs in the most abundant bacteria, *F. sanfranciscensis*, and found they were present in all sourdoughs. By contrast, the ASVs of the dominant sourdough yeast species (*S. cerevisiae, K. bulderi* and *K. humilis*) were generally specific to a single sourdough (Figure 6). However, some ASVs were found in several sourdoughs. Sourdoughs from bakers B12, B15, B26 and B63 shared one *K. bulderi* ASV. Sourdoughs from bakers B04, B17, B31, B56 and B58 shared one to three *K. humilis* ASVs. Sourdoughs from bakers B29, B55 and B74 shared one to two *S. cerevisiae* ASVs and those from bakers B01, B16, B17 and B32 shared another ASV. Bakers who shared a yeast ASV generally belonged to the same bakery practices cluster. The group of three bakers who shared one or two *S. cerevisiae* belonged to cluster 2 (corresponding to artisanal bakery practices) while three of the four bakers who shared another *S. cerevisiae* ASV belonged to cluster 1 (corresponding to farmer-baker practices). Three of the five bakers who shared at least one *K. humilis* ASV belonged to cluster 2, but the two others, who belonged to cluster 1, shared more ASVs than with the three others (Figure 5). An evaluation of the association between sourdough fungal community dissimilarity and geographical distances did not reveal any significant correlation (mantel z = 363535.1, P = 0.547). The only link that could be made from the data on sourdough exchanges concerned farmer-baker B15, who shared a *K. bulderi* ASV with farmer-baker B12, and started his sourdough using B12 sourdough.

**Figure 6-.**
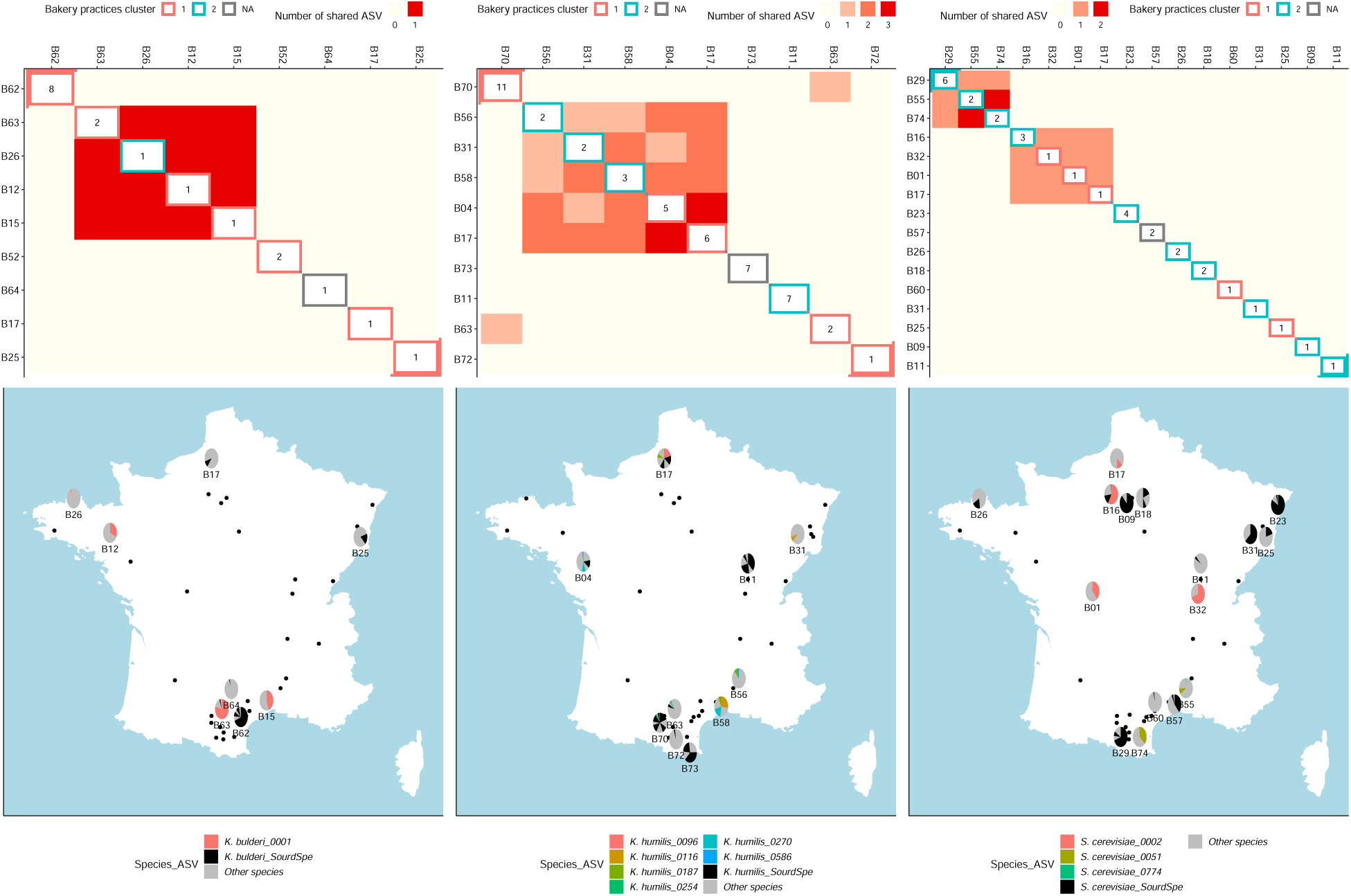
Sourdoughs sharing *K. bulderi, K. humilis* and *S. cerevisiae* ASVs. Top, the heatmaps show the number of shared ASV between sourdoughs, each tile being colored according to the number of shared ASV. In the diagonal, the number of ASVs of the considered species in each sourdough are displayed, and the tiles are underlined according to the cluster of bread-making practices (1 = farmer-baker and 2 = artisan-baker). At the bottom, the maps of France show the locations of each baker. Bakers are represented by a point when the species considered was not detected in their sourdough, and in the other case the pie charts show the composition of their sourdoughs. ASVs that are shared between at least two different sourdoughs are colored and their identifiers displayed in the legend, while the ASVs of species considered to be specific to one sourdough are represented in black (SourdSpe in the legend), while ASVs from other species are in grey.

## 4 Discussion

The composition of the sourdough microbiota was consistent with previous studies on sourdough. The mean LAB to yeasts ratio in sourdoughs was 65.4, which is within the same range as that reported by other studies (Zhang et al., 2011; Lhomme et al., 2015; Arici et al., 2017; Fraberger et al., 2020). As previously detected in French sourdoughs, *F. sanfranscisensis* was the most frequently encountered bacterial species. *S. cerevisiae, K. humilis* and *K. bulderi* were the most frequently encountered sourdough dominant yeast species (Michel et al., 2016; Urien et al., 2019; Lhomme et al., 2015). Moreover, *K. bulderi* was associated with farmer-baker practices, as previously reported by Michel et al. (2019). Surprisingly, *Saccharomycetales* accounted for fewer than 5% of the reads in ten sourdough samples, yet a typical yeast density and morphology was observed in almost all of these samples. This may have reflected biases in the metabarcoding analysis (Loos and Nijland, 2020). DNA could have been poorly extracted or amplified, thus leading to a low number of reads. The reads might also not have passed the quality filtering or merging steps in the bioinformatics analysis, particularly if the ITS region was too long. This is a limitation of the dada2 software, where reads that are too long to be merged are lost. However, this does not concern the ITS database, as in this case the ASV would have been found but not assigned to a species.

### 4.1 Flour-associated species were mainly plant-associated microorganisms

The microbiotas of the flours mainly comprised plant-associated microorganisms. Several filamentous fungi known to be cereal pathogens, and notably *Alternaria* and *Mycosphaerella* species, were detected. Similarly, several bacterial genera such as *Pseudomonas* and *Pantoea* were found. Many species in these genera are plant pathogens or plant-associated species (Dutkiewicz et al., 2016; Preston, 2004).

Most of the species that we detected in flour during this study had been mentioned in previous studies on wheat seed microbiotas (Kuzniar et al., 2020; Rozhkova et al., 2021; Minervini et al., 2018b). They had also been mentioned in studies describing flour microbiota, and the results were in accordance with those of De Angelis et al. (2019) who compared the microbiotas of soft and *durum* wheat flour using culture independent methods. Minervini et al. (2018a) analyzed the microbiotas of three different flours, and found the species *F. sanfranciscensis* in every sample (4% of all the strains isolated from the flour). This was higher than what we found, and could have been related to bias affecting the culture independent analyses, where rare species can go undetected.

Surprisingly, the filamentous fungi plant-associated pathogens detected in flour were also detected in sourdoughs. However, on average they accounted for 54% of the reads (sd = 30%) in sourdough and 92% (sd = 9.3%) in flour, suggesting that filamentous fungi die in the acidic environment of sourdough and/or are poor competitors with yeasts in this environment. To our knowledge, they have never been detected alive in sourdough, even though they are able to grow on the media classically used to enumerate yeasts(Me and Melvydas, 2007). The presence of their DNA in sourdough suggested that this was partly protected in this environment, possibly thanks to their cell wall structure. The high proportion of these fungi in sourdough may also be related to bias affecting DNA extraction and amplification.

Unlike filamentous fungi, the common plant bacteria *Pantoea* and *Pseudomonas* were not detected in sourdoughs, suggesting they did not survive in the sourdough ecosystem and that their DNA was degraded. This is highly probable as *Pseudomonas* species generally do not survive at a low pH.

### 4.2 LAB found in flour were typical of the first stage of sourdough preparation

As well as plant pathogens, the flour microbiotas contained several LAB genera : *Lactococcus, Pediococcus* or *Weisella*. They had all been detected previously at the first stages of new sourdough preparation (Bessmeltseva et al., 2014), before being replaced by other LAB species, generally *F. sanfranciscensis*. The bacterial species present during the early stages of sourdough preparation may therefore arise from the flour. They do not benefit from a priority effect, and the succession of microbial communities during sourdough initiation does not follow the pattern of the community monopolization hypothesis (Nadeau et al., 2021), where an early arriving species can adapt to the environment and gain a competitive advantage over previously better adapted species, thereby altering the community assembly.

### 4.3 LAB present in flour did not develop in mature sourdough

However, our results showed that mature sourdoughs did not contain the same LAB as those provided by the flour. *F. sanfranciscensis*, which is the most frequently encountered LAB species in sourdough, was almost never found in flour. The most abundant *F. sanfranciscensis* ASV in sourdoughs, which is shared across all the French sourdoughs studied, was never detected in flour samples. It may have been absent from the flour, or present at very low levels, and was thus not detected by the metabarcoding analysis. Because the microbial counts were very low in flour (around 10^3^ CFU/g), the species may not have been detected. Nevertheless, rare *F. sanfranciscensis* ASVs were detected in five flour samples, but these flour ASVs were only detected in one case in the related sourdough. Because the V3-V4 region of 16S rRNA displays low intraspecies diversity, we can consider that the different bacterial ASVs corresponded to different strains. Therefore, the *F. sanfranciscensis* strains found in flour did not appear to develop in an established sourdough. This contradicts the findings of (Minervini et al., 2018a), who determined the same strains of *F. sanfranciscensis* in flour and sourdough in three bakeries.

### 4.4 Flour as the source of sourdough microorganisms

Two hypotheses could be advanced concerning evolution of the sourdough population of *F. sanfranciscensis*. On the one hand, the bacterial population may have come from an ancestral flour population and subsequently evolved. On the other hand, the sourdough and flour bacterial populations could have separate ancestral origins, with the sourdough population arising from a a source other than flour, such as the baker’s hands, bakery equipment, or insects, etc. Further investigation of the intraspecific diversity of *F. sanfranciscensis* is necessary to shed light on its origin and evolutionary dynamics.

During this study, none of the yeast species usually found in sourdough was detected in flour samples, so the sourdough yeasts did not appear to have come from the flour. The preferential occurrence of *K. bulderi* in sourdoughs made by farmer-bakers or *K. humilis* in artisanal sourdoughs could not be explained by the different flour supply chains. This finding is in agreement with previous studies which showed that the species composition of sourdough yeasts depended more on the bakery house than on the cereal flour species used (Minervini et al., 2015; Comasio et al., 2020).

### 4.5 Yeast dispersal between sourdoughs

The exchange or gifting of sourdoughs between bakers can lead to yeast dispersal, as was found between bakers B15 and B12 who were regularly in contact and exchanged their sourdoughs. However, this practice is not common, as bakers prefer to develop their own sourdough when they lose one. Finding the same sourdough yeast species in dough from several farmer-bakers could be explained by the development of networks of bakers who meet to share their knowledge and skills, and yeast dispersal may be promoted through handshakes. Student bakers traveling between different bakeries may also be a vector for dispersal. Bakers belonging to the same bakery practices cluster (artisanal or farmer-baker) tended to share more ASVs than with bakers from the other cluster (see Figure 6). However, the number of bakers sharing yeast ASVs was quite low : four, six, and eight bakers shared at least one *K. bulderi, K. humilis* and *S. cerevisiae* ASV respectively, so we were not able to perform a robust statistical analysis.

A population genomic analysis of *K. bulderi, K. humilis* and *S. cerevisiae* from sourdoughs would shed more light on the relative impact of gene flow and selection on the evolution of these sourdough yeasts. The genomes of *K. bulderi* and *K. humilis* were released recently (BioProject accession number PRJEB44438 in the NCBI BioProject database, https://www.ncbi.nlm.nih.gov/bioproject/) and this will enable the conduct of these studies.

In conclusion, this evaluation of the bacterial and fungal composition of flour and sourdough showed that their microbiotas overlapped little. Flour did not appear to act as a vector for the dispersal for sourdough yeasts, but might be a vector for the dispersal of sourdough LAB. However, the LAB carried by the flour were not able to develop in a mature sourdough.

## Acknowledgments

We would like to thank Sylvain Santoni and Audrey Weber for the Illumina sequencing and their valuable advices. This work was partly supported by a grant from the Fondation de France (Gluten : mythe ou réalité ?). Authors thank Dominique DESCLAUX, Kristel MOINET, and all the bakers and farmer-bakers that have shared their sourdough, flour and knowledges.

## Data accessibility

Raw data and scripts are available at :

https://data.inrae.fr/dataset.xhtml?persistentId=doi:10.15454/DF0BRL

## Conflict of interest

The authors declare no conflict of interest

## Author contributions

## Notes

https://data.inrae.fr/dataset.xhtml?persistentId=doi:10.15454/DF0BRL

### Competing Interest Statement

The authors have declared no competing interest.

