## Supplemental Table 1 for "Microbial community dispersal in sourdough"

Table S1 : Detailed list of the primers used for the metabarcoding analysis.

| Genomic region | Primer name | Primer sequence |
|---|---|---|
| 16S V3-V4 | Illu4_16SV3F | TCGTCGGCAGCGTCAGATGTGTATAAGAGACAGNNNNTACGGRAGGCAGCAG |
| 16S V3-V4 | Illu6_16SV3F | TCGTCGGCAGCGTCAGATGTGTATAAGAGACAGNNNNDVTACGGRAGGCAGCAG |
| 16S V3-V4 | Illu8_16SV3F | TCGTCGGCAGCGTCAGATGTGTATAAGAGACAGNNNNNNRMTACGGRAGGCAGCAG |
| 16S V3-V4 | Illu4_16SV4R | GTCTCGTGGGCTCGGAGATGTGTATAAGAGACAGNNNNTACCAGGGTATCTAATCCT |
| 16S V3-V4 | Illu5_16SV4R | GTCTCGTGGGCTCGGAGATGTGTATAAGAGACAGNNNNHTACCAGGGTATCTAATCCT |
| 16S V3-V4 | Illu6_16SV4R | GTCTCGTGGGCTCGGAGATGTGTATAAGAGACAGNNNNNWTACCAGGGTATCTAATCCT |
| ITS1 | Illu4_ITS1F | TCGTCGGCAGCGTCAGATGTGTATAAGAGACAGNNNNCTTGGTCATTTAGAGGAAGTAA |
| ITS1 | Illu6_ITS1F | TCGTCGGCAGCGTCAGATGTGTATAAGAGACAGNNNNDVCTTGGTCATTTAGAGGAAGTAA |
| ITS1 | Illu8_ITS1F | TCGTCGGCAGCGTCAGATGTGTATAAGAGACAGNNNNNNRMCTTGGTCATTTAGAGGAAGTAA |
| ITS1 | Illu4_ITS2R | GTCTCGTGGGCTCGGAGATGTGTATAAGAGACAGNNNNGCTGCGTTCTTCATCGATGC |
| ITS1 | Illu5_ITS2R | GTCTCGTGGGCTCGGAGATGTGTATAAGAGACAGNNNNHGCTGCGTTCTTCATCGATGC |
| ITS1 | Illu6_ITS2R | GTCTCGTGGGCTCGGAGATGTGTATAAGAGACAGNNNNNWGCTGCGTTCTTCATCGATGC |
